# clusTCR: a Python interface for rapid clustering of large sets of CDR3 sequences

**DOI:** 10.1101/2021.02.22.432291

**Authors:** Sebastiaan Valkiers, Max Van Houcke, Kris Laukens, Pieter Meysman

## Abstract

The T-cell receptor (TCR) determines the specificity of a T-cell towards an epitope. As of yet, the rules for antigen recognition remain largely undetermined. Current methods for grouping TCRs according to their epitope specificity remain limited in performance and scalability. Multiple methodologies have been developed, but all of them fail to efficiently cluster large data sets exceeding 1 million sequences. To account for this limitation, we developed clusTCR, a rapid TCR clustering alternative that efficiently scales up to millions of CDR3 amino acid sequences. Benchmarking comparisons revealed similar accuracy of clusTCR with other TCR clustering methods. clusTCR offers a drastic improvement in clustering speed, which allows clustering of millions of TCR sequences in just a few minutes through efficient similarity searching and sequence hashing.

clusTCR was written in Python 3. It is available as an anaconda package (https://anaconda.org/svalkiers/clustcr) and on github (https://github.com/svalkiers/clusTCR).

## Introduction

T cells constitute a key component of the adaptive immune system and are one of the primary determinants of distinguishing self from non-self. The T-cell receptor (TCR) is responsible for the recognition of peptide antigens presented by the major histocompatibility complex (MHC). The TCR complex is a heterodimer consisting of two distinct chains (*α* and *β*) that both contribute to the recognition of the cognate antigen and is expressed on the T-cell surface. High-throughput targeted sequencing technology enables sequencing of the unique and diverse TCR *α* and/or *β* nucleotide sequences in a sample, allowing quantitative mapping of the immune receptor repertoire. One of the major goals of quantitative immunology is the identification of groups of T cells with common specificity towards an antigen. Exactly determining a TCR’s epitope-specificity requires knowledge about the epitope and demands for time-consuming *in vitro* experiments such as MHC multimer assays (1). An alternative way of characterising specificity groups is unsupervised clustering of TCR sequences. This does not require prior knowledge of specific epitopes and allows interrogation of complete repertoire data sets by searching for sequentially similar TCRs. It has been illustrated previously that TCRs sharing similar CDR3 sequences often target the same epitope (2). Hence, an effective approach for TCR clustering is determining the sequence-based similarity within a set of TCR sequences. However, in spite of being a powerful approach to drastically reduce the repertoire complexity, sequencebased clustering has various complications. The most pronounced bottleneck of clustering sequence data is the scalability of pairwise distance calculations. Calculating pairwise distances scales quadratically with the number of input sequences (*O*(*n*^2^)). Current methods for TCR clustering rely on a pairwise distance matrix to determine the clusters. Consequently, these algorithms have limited applicability towards extremely large RepSeq data sets. With clusTCR, we created a clustering procedure that can efficiently classify large sets of CDR3 sequences into specificity groups by drastically limiting the number of required pairwise comparisons.

## Methods

### Data

#### Epitope-labeled data

We used TCR sequences with known epitope specificity to benchmark the clustering quality of our method. For this we downloaded all human TRB sequences from the VDJdb (3). This provided us with a list of 28,576 unique CDR3 sequences. Each TCR-epitope pair in VDJdb is annotated with a quality score, which ranges from 0 to 3 (with 3 indicating the highest quality database entries). The VDJdb quality score reflects the confidence of the antigen specificity annotation. The two subsamples contained sequences that satisfied quality scores of ≥ 1 and ≥ 2. The subsamples consisted of 2851 and 495 unique CDR3 sequences respectively.

#### Unlabeled data

Due to the limited availability of epitope-labeled TCR sequencing data, unlabeled TCR repertoire data was used to evaluate the speed of clusTCR and other TCR/CDR3 clustering methods. The data set by Emerson et al. (2017) (4) contains a large amount of unique CDR3 sequences and is therefore well suited for performance benchmarking. The data was downloaded from the immuneACCESS database. From this data set we randomly sampled a fixed number of unique sequences to construct ‘metarepertoires’. These metarepertoires may therefore contain CDR3 sequences that originate from different repertoire samples. For the performance benchmarking, different metarepertoires were sampled (see table S1).

### clusTCR workflow

#### Overview

clusTCR is a two-step clustering algorithm that combines the speed of the Faiss library, combined with the accuracy of the Markov clustering algorithm (MCL). During the first clustering step we use Faiss’ efficient K-means implementation to rapidly subdivide data sets of CDR3 sequences into superclusters. In the next step, MCL is applied on each individual supercluster to identify groups of epitope-specific CDR3 sequences.

#### Sequence vectorization

First, CDR3 amino acid sequences are converted to *n*-dimensional vectors. *n* must be predefined and will be the same for every sequence. Therefore, the default number of dimensions is equal to the length of the longest sequence in the input data. The vectorized sequences reflect the physicochemical properties of the amino acids. Different combinations of physicochemical properties were tested and the resulting clustering quality and speed were evaluated (for an exhaustive list of the combinations, see fig. 2). All vectors are stored in one large vector matrix with shape (*n, m*), where *n* is the size of a single vectorized sequence and *m* is the total number of sequences in the input data.

#### Computing superclusters

Roughly dividing the sequence data into groups of approximate neighbours drastically speeds up the total clustering process by minimizing the number of direct comparisons between sequences. clusTCR computes a number of superclusters *s.* This number *s* corresponds to *m/θ,* the total number of sequences in the data set (*m*) divided by the average number of sequence in a supercluster (*θ*). The latter is set to 5000 by default. To retrieve the superclusters, we use the rapid K-means implementation from the Facebook artificial intelligence similarity search (Faiss) Python library (5). First, an index is constructed (‘trained’), which will be used to efficiently search large sets of CDR3 sequences. During this step *m/*5000 centroids are computed. Next, we use Faiss’ efficient K-means implementation to assign each vector to its most similar centroid (as defined by the Euclidean or L2 distance between the vector the centroids).

#### Sequence hashing to speed up graph construction

Networks are effective structures for representing relationships between objects or processes, such as biomolecular interactions. Likewise, networks can be used to represent the sequence similarity landscape of CDR3 amino acid sequences. These similarity networks are undirected graphs *G*(*V,E*), consisting of a set of nodes or vertices (*V*) represented by the CDR3 sequences, which are connected by a set of edges (*E*) representing (dis)similarity between nodes. In the case of amino acid sequences, dissimilarity can be described by an edit distance. Here, edit distance was defined as the Hamming distance (HD) between two strings. Given two strings *u* and *v,* the HD between them (*d*(*u,v*)) is defined as the number of positions where *u* and *v* differ. Additionally, HD implicitly assumes that both strings have an identical length.

Thus HD only allows substitutions, and no deletions or insertions. The graph describes the adjacency matrix *A* of pairwise HDs between the CDR3 amino acid sequences:

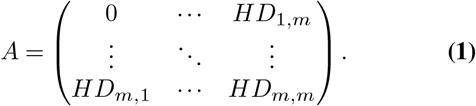

HD = 1 was chosen as a criterion for drawing an edge between two sequences. Hence *A* will be a binary matrix that can be compressed into a sparse matrix format for efficient memory allocation.

To determine all pairs of CDR3 sequences with HD = 1, a simple hash function was used to convert each sequence into hashes that contain either the odd or even positions of the original sequence. By doing this, only sequences that are assigned to the same hash need to be compared against each other. This drastically reduces the total amount of pairwise comparisons. This hashing method correctly identifies all sequence pairs with a HD of 1, while being much faster than brute-force comparison methods.

#### Graph clustering

For each individual supercluster, a graph is constructed from its CDR3 sequences. The goal is to resolve clusters of sequences with high similarity (~ epitope-specificity). MCL is particularly appropriate for this task (6). MCL is a graph clustering algorithm that identifies dense network substructures in a graph or network by simulating stochastic flow. The algorithm performs a random walk on the graph, which is calculated using Markov chains. MCL performs two operations on the stochastic matrix: *expansion* and *inflation.* Expansion is taking the power of the stochastic matrix using the normal matrix product. This determines how much flow is allowed between different regions of the graph. This effect is further exaggerated by the inflation parameter, taking entrywise powers of the ’expanded’ matrix, followed by a rescaling step so that the matrix elements correspond to probability values (i.e. making the matrix stochastic again). Inflation further strengthens current between (already) strong neighbours, while at the same time weakening current between weak or distant neighbours. This process of alternating between expansion and inflation operations is repeated until the graph is partitioned into individual substructures (while paths between substructures no longer exist). We evaluated different thresholds of the expansion and inflation parameters, as illustrated in fig. S2.

clusTCR applies the Python 3 implementation of MCL (see github). To further increase performance, clusTCR also applies multiprocessing to parallelise MCL for different superclusters at the same time.

#### GPU support & batch clustering

To enhance the clustering performance for extremely large data sets, clusTCR offers GPU support for CUDA-compatible GPUs and implements batch clustering to prevent memory overflow. GPU support is provided for the calculation of superclusters. To allow the use of the GPU, the user must ensure having installed the required cudatoolkit (installation instructions available in the software documentation). For extremely large data sets that do not regularly fit into RAM, clusTCR provides a batch clustering functionality. Here, cluster centroids are computed from a subset of the complete data set. We recommend a subset of size 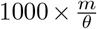, where *m* is the size of the complete data set to be clustered, and *θ* is the average size of the superclusters (a standard parameter of clusTCR, which is 5,000 by default). Next, vectors are assigned to the closest centroid, in batches. During this step, intermediate results are stored on disk, with a file for each supercluster. Next, the second clustering step is also performed in batches. Using a generator function, data is loaded and clustered batch per batch. To optimize this process further, multiple superclusters are loaded at once and clustered using multiprocessing.

### Clustering evaluation

#### Evaluation metrics

Evaluating the quality of clustering results is not a trivial task, even when ground truth labels (in this case, the epitope to which the TCR sequence is specific) are available. Important factors to consider when evaluating TCR clustering results are the coverage of the clustering method, the extent to which epitope-specific sequences are clustered together and how consistent this process is. The following metrics for the evaluation clustering quality were set out to evaluate these criteria: retention, purity and consistency. Although each metric has its limitations, collectively they offer a good means of evaluating an unsupervised clustering approach like clusTCR.

#### Retention

Not all input sequences end up in the final clustering results. clusTCR only considers those sequences that share at least one neighbour with a hamming distance (HD) of 1. We define retention as the number of TCR/CDR3 sequences *s* that participate to any cluster *c* divided by the size of the data set.

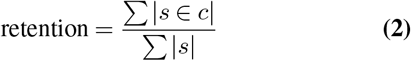

#### Purity

Purity is a simple and interpretable metric to describe the extent to which clusters contain a single epitope. Purity is calculated as the fraction of sequences within one cluster targeting the same epitope. For each cluster *c*, we count the number of sequences *s* specific for the most common epitope *γ*, sum the values and divide them by the total number of sequences in any cluster. Formally, purity is computed as

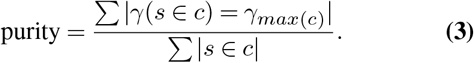

Since clustering methods that return very small clusters (i.e. cluster size = 2-3) automatically achieve high purity, we also calculated the fraction of clusters with purity > 90% as an alternative metric for clustering quality. Thereby, more weight is assigned to larger clusters.

#### Consistency

Consistency describes the fraction of epitope-specific CDR3 sequences that are assigned to a single cluster. In the case of having multiple epitopes assigned to one cluster, the largest epitope (i.e. with the most sequences specific to it) is given preference. Also, if two or more clusters contain sequences specific for that epitope, the cluster containing the most epitope-specific CDR3 sequences is assigned as the true cluster. Formally, we computed consistency as:

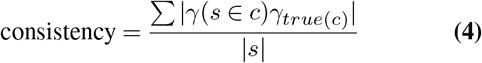

### Benchmarking clusTCR

#### TCR clustering algorithms

To benchmark the performance of clusTCR, it was compared against existing methods for clustering TCR sequences. For each method, the performance and clustering accuracy were evaluated using the same data. Quality benchmarking was performed using different subsets of the VDJdb, while the performance of the different algorithms was evaluated on artificially large metarepertoires of different sizes (see table). Note that all methods, excluding clusTCR, additionally require V gene information, hence this was also provided.

- **iSMART:** The code for the iSMART algorithm was downloaded from github (link). Although iSMART implements parallel CPU processing, we encountered fatal kernel errors when using this feature. Therefore, iSMART was used at 1 CPU. Additionally, a kernel error appeared when trying to cluster *>* 200K sequences.
- **GLIPH2:** We downloaded the .centos executable version of GLIPH2 from the website and installed it in the clusTCR repository. Each GLIPH2 input TCR included a CDR3 sequence, V gene, subject id and frequency. We wrote a python wrapper that provided GLIPH2 with the appropriate data and executed the algorithm.
- **tcrdist3 (+ DBSCAN):** Pairwise distances between TCR sequences were calculated using the *tcrdist3* python package (github). Since *tcrdist3* does not offer clusteringing procedures, we used density-based spatial clustering of applications with noise (DBSCAN) from the python scikit learn library, which was previously shown to be an appropriate method for clustering CDR3 sequences [(2)]. Another reason why we used DBSCAN is that the algorithm, like MCL, does not require to define the number of clusters in advance (unlike e.g. K-means). The minimum amount of items per cluster was set to two. We used different similarity thresholds, thereby evaluating the trade-off between retention and purity/consistency. Ultimately, the threshold was set to 18. This threshold best reflected the outcome of the other clustering methods used for benchmarking.

### Downstream cluster analysis

#### Cluster features

To allow downstream machine learning applications of the clustering results, clusters were represented numerically by calculating a feature matrix describing multiple properties of the CDR3 amino acid sequences in a cluster.

- **CDR3 length:** Length of CDR3 sequences (number of amino acids) in the cluster. Since clusTCR handles with an exact similarity criterion of HD = 1, all sequences within one cluster have an equal length.
- **Cluster size:** The number of sequences in a cluster.
- **Cluster entropy:** Cluster entropy describes the positional variation within a cluster. It is calculated as the average information content within a cluster by determining the Shannon entropy at each position (excluding positions 0 and −1) and taking the average across all positions. Additionally, small-sample correction is applied to account for differences in cluster size (7). Since all sequences within a single cluster have an equal size, multiple sequence alignment is not necessary. Formally, we compute the average information content *R* of a cluster:

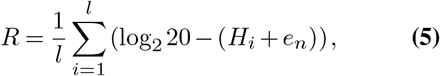

where *l* equals the length of the sequences in the cluster and *n* represents the number of sequences in the cluster. *H_i_* is the Shannon entropy at a given position *i*:

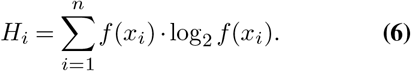

*e_n_* is a correction factor that accounts for differences in cluster size. It is calculated as folows:

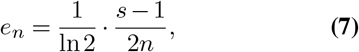

where *s* is the size of the alphabet. Because we are handling amino acid sequences, *s =* 20.
- **Physicochemical features:** clusTCR calculates the average and variance of following physicochemical features basicity, hydrophobicity, helicity and mutation stability. Physicochemical features provide a functional encoding of the sequences within a cluster.
- **Generation probability:** Generation probability (*P_gen_*) is the probability by which a TCR or CDR3 sequence is generated through the V(D)J recombination process. This probability is estimated through stochastic modelling of V(D)J recombination. *P_gen_* values were calculated using the Optimized Likelihood estimate of immunoGlobulin Amino-acid sequences (OLGA) algorithm (8).

#### Predicting cluster quality

To analyse clusTCR’s clustering output, a binary classification system was constructed that predicted the whether a cluster is of high quality or not. Cluster quality was here defined as having at least 0.90 purity. Labels and features were generated by performing clusTCR two-step clustering on all unique CDR3 sequences from the human TRB sequences in the VDJdb. Purity of each individual cluster was calculated and these purity values were discretized according to the defined purity cut-off. Clusters with purity > 0.90 were assigned 1, every cluster with lower purity was assigned 0. Next, cluster features were calculated according to the procedure described in the previous section. We used Python’s scikit learn library to train a random forest classifier consisting of 100 decision trees using the cluster features and labels. The classifier was evaluated through 10-fold stratified cross-validation. We used the receiver operating characteristic (ROC) and the area under the ROC (AUC) to express the performance of this classification model.

## Results

### Choosing hyperparameters

clusTCR is a two-step clustering approach that combines the speed of the Faiss library with the accuracy of MCL (fig. 1). The first step drastically scales down the search space by subdividing the data into large sets of sequences sharing some degree of similarity. We refer to these groups as superclusters. Faiss’ efficient K-means implementation is used to generate these superclusters. Because Faiss efficiently handles vectors, sequences were first encoded as numerical vectors reflecting their physicochemical properties. Next, *n* centroids were predefined to which the vectorized sequences are assigned. The number of centroids *n* is a function of the average supercluster size. Although the limited availability of epitope-labeled CDR3 data prevents us from exactly determining an optimal supercluster size, 5000 gave consistent results while maintaining high performance (fig. S1). Next, the effect of sequence encoding was assessed by evaluating different combinations of features. No specific combination of physicochemical properties was found to significantly outperform others (as determined by cluster quality evaluation, fig. 2A,B). Combinations including isoelectric point (pI) took significantly longer to compute and were therefore excluded (fig. 2C). During benchmarking of clusTCR, a combination of mutation stability and amino acid z-scores (derived from quantitative structure–activity relationship studies (9)) was used.

**Fig. 1.**
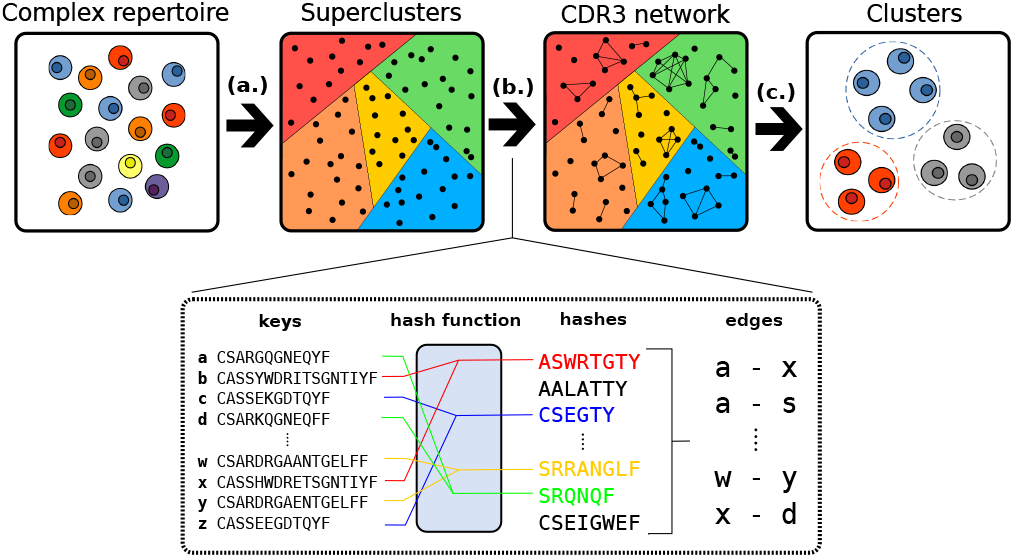
Workflow of clusTCR. **a.** Sequences are first roughly categorized into superclusters through an efficient nearest-neighbor search. **b.** Within each supercluster, a hash function is applied to sort sequences. Sequence pairs with a maximum edit distance of 1 are selected from each hash. These sequence pairs are used to construct a graph. **c.** MCL is used to find dense network substructures.

**Fig. 2.**
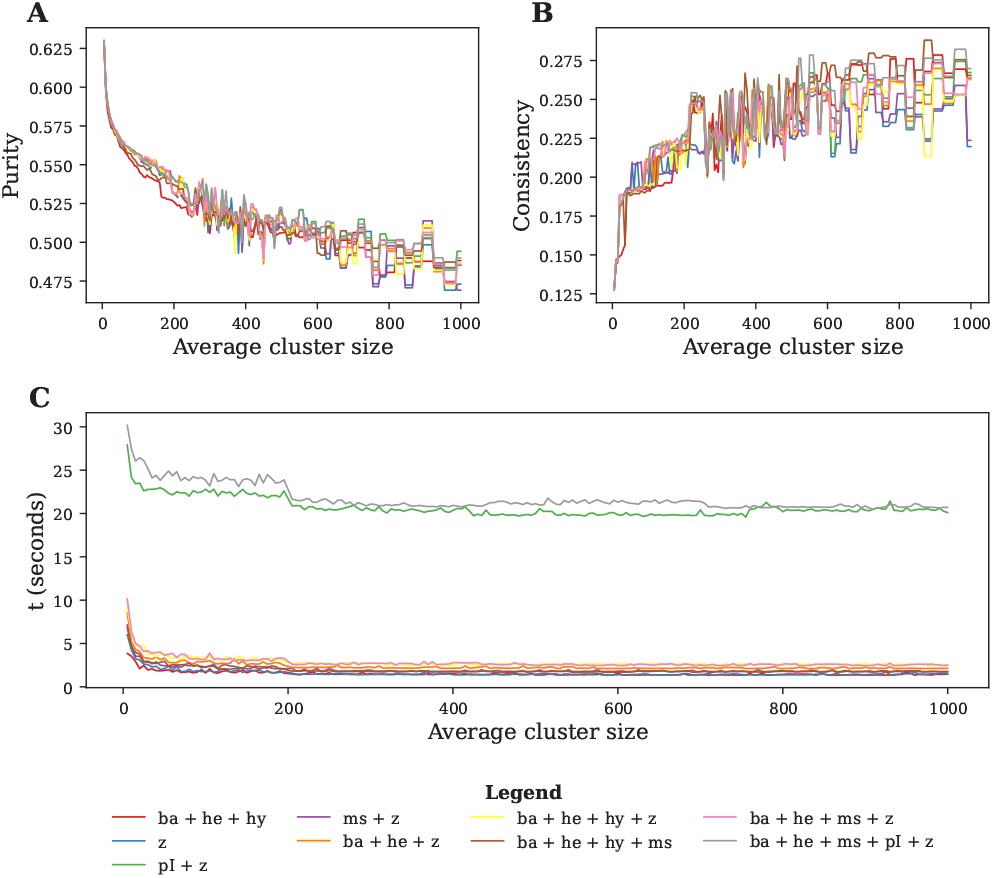
Sequence vectorization for efficient K-means clustering. Different combinations of features were used to numerically encode CDR3 sequences to allow vector clustering with Faiss’ K-means implementation. **A.** Influence of different features on cluster purity. **B.** Influence of different features on cluster consistency. **C.** Elapsed time for computing different numerical vectors. Combinations including isoelectric point (pI) take significantly longer to compute. Abbreviations: ba, basicity; he, helicity; hy, hydrophobicity; ms, mutation stability; pI, isoelectric point; z, amino acid z-scores.

In the second step, clusTCR reclusters each individual supercluster to accurately identify specificity groups within. Adaptive immune receptor repertoires can be represented as graphs in which nodes represent the sequences and the edges represented similarity between sequences (10). To create this graph, clusTCR uses efficient sequence hashing to determine each pair of sequences with an exact Hamming distance of 1. Next, it uses the corresponding similarity-grouped graph to identify potential epitope-specific clusters. Evaluating the graph structure allows the interrogation of sequence-based relationships in the repertoire because similar sequences will share edges within the graph. To this end, clusTCR applies MCL for the identification of dense network substructures (11), representing dense groups of CDR3 sequences with similar sequential characteristics. MCL simulates stochastic flow inside the graph by alternating between two operations performed on the stochastic matrix (this matrix describes the probabilities of visiting a specific node): expansion and inflation. Hereby, MCL identifies dense network substructures where flow is high. These network substructures represent the clusters in clusTCR’s output. Different combinations for expansion and inflation were tested (fig. S2). Increasing the expansion parameter typically resulted in an overall decrease in purity, but an increase in consistency and the fraction of clusters with purity > 0.90. Increasing the inflation parameter had an opposite effect. Individual application of MCL allows efficient and accurate clustering of relatively small sets of CDR3 sequences (up to 50,000). Combined with the first step, the clustering can be made efficient for any TCR dataset.

### Benchmarking

In order to effectively validate clusTCR, we compared it against existing TCR/CDR3 clustering approaches. These approaches included iSMART (12), GLIPH2 (13) and tcrdist3 (14). Both iSMART and GLIPH2 have illustrated evidence for being suitable approaches for clustering TCR sequences. tcrdist was developed as a method for defining (dis)similarity between TCR sequences. The latter does not directly generate clusters, rather a pairwise distance matrix. Consequently, we used DBSCAN to perform clustering based on the pairwise distances determined by tcrdist3. The DBSCAN algorithm requires a predefined distance threshold to cluster a set of items. We evaluated a range of similarity thresholds (fig. S3). A similarity threshold of 18 was selected, as this reflected the clustering results of the other algorithms. Indeed, this has previously been shown to be a representative similarity threshold for tcrdist-based distance matrices (2).

The ability of different clustering methods to group CDR3/TCR sequences with common epitope-specificity was tested using a data set of sequences with known epitope-specificity. From this data set we selected two subsets of database entries with a VDJdb score of > 1 and > 2 respectively. This score reflects the confidence of the antigen specificity annotation. We evaluated each clustering method on the unfiltered set of human TRBs in the VDJdb, and the two high-quality subsets. On the smallest subset (495 sequences), both GLIPH2 and clusTCR achieved near perfect purity (0.99 and 0.98 respectively), outperforming iSMART (0.91) and tcrdist3 (0.80) on this metric (fig. S4A). GLIPH2, clusTCR and iSMART achieved achieved a purity of ~ 0.88 on the second subset (2,851) sequences. tcrdist3 performed slightly worse, achieving an overall purity of 0.80. When clustering the complete data set (28,576 sequences), both GLIPH2 and iSMART (0.66 both) achieve slightly higher purity compared to clusTCR (0.60). Since smaller clusters automatically achieve higher purity, we used the fraction of clusters with purity > 0.90 as an additional metric to account for this limitation. Evaluating clustering results with this metric revealed equal performance of all methods at every confidence level (S4B). However, tcrdist3-based clustering underperformed as compared to other methods (66% versus ≥ 90%). There does exist a trade-off for all TCR clustering methods between the number of clusters and the size or quality of the clusters. As illustrated, clusTCR covers only 20-25% of the input sequences of the ground truth data set. This is lower compared to GLIPH2 and tcrdist3, but higher than iSMART (fig. S4C). However this increased cluster retention comes at the cost of clustering consistency (S4D). In this regard, clusTCR comes out on top along with iSMART. Collectively, these results suggest that there is no single approach that clearly outperforms all others with regards to the final cluster quality.

We then evaluated the speed of clusTCR as compared to existing TCR clustering approaches (fig. 3). Metarepertoires of various sizes (ranging from 5,000 to 1,000,000 unique sequences) were randomly sampled from a large immunosequencing data set by Emerson et al. (2017) (4). For small data sets (5,000 to 100,000 sequences), clusTCR outperforms all other methods by at least 2.5-(5,000 sequences) to 23.0-fold (100,000 sequences). From our testing, both iSMART and tcrdist3 failed to cluster medium-sized data sets of > 200,000 sequences. At very large data set sizes, clusTCR achieves up to 50× speed improvement compared to GLIPH2. In absolute numbers, clusTCR successfully clusters 1 million sequences in 4 minutes (±30 seconds). GLIPH2 clustered the same amount of sequences in approximately 2 hours and 45 minutes (±11 minutes).

**Fig.3.**
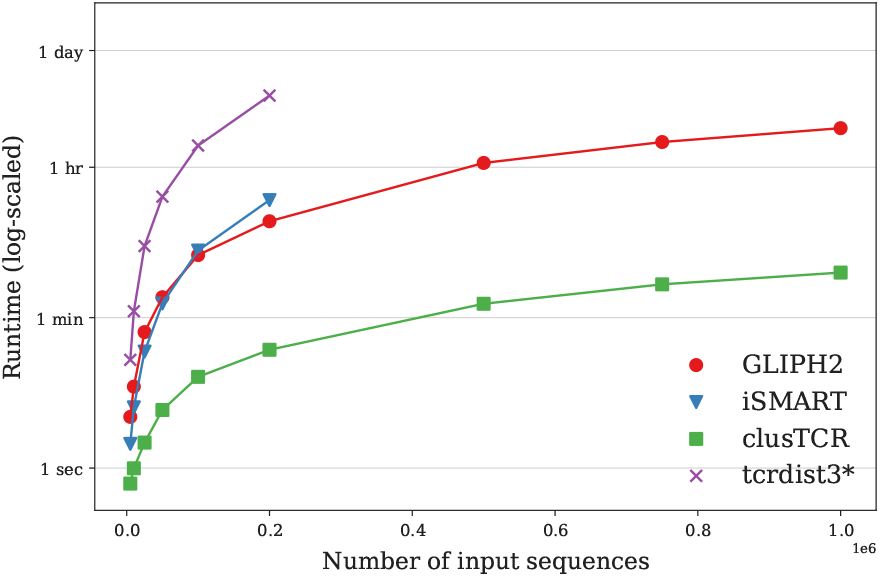
PerformanceofdifferentTCR/CDR3 clustering methods. At 10^6^ sequences, clusTCR provides a > 50× speed improvement (with an Intel(R) Core(TM) i7- 10875H CPU @ 2.30GHz, using 8 CPUs) over other methods. *: The runtime of tcrdist3 only includes the computation of pairwise distances, not clustering.

### Downstream clustering analysis

After determining clusters of CDR3 sequences, we calculated a range of features for each cluster. By numerically representing the clustering results, we can unlock downstream machine learning applications for repertoire classification or cluster enrichment analyses. Calculated features included cluster entropy, physicochemical properties and generation probability. Using these features, a classification model was constructed that differentiates between high- and low-quality clusters. High-quality clusters were defined as clusters with purity > 90%. A random forest classifier, consisting of 100 decision trees, was fitted with the cluster features and their associated, discretized labels (1: purity > 90%, 0: < 90%). The classification performance of the model was evaluated through 10-fold stratified cross-validation. An average AUC of 0.84 (±0.07) was achieved (fig. S5A). Evaluation of the feature importances revealed a significant contribution of generation probability to the prediction of clustering quality (fig. S5B). We propose that this model can aid in filtering out insignificant, low-quality clusters.

## Discussion

We have developed clusTCR, a rapid algorithm for clustering large sets of CDR3 sequences. clusTCR was benchmarked by comparing it to three other naive clustering methods for TCRs or CDR3 sequences: GLIPH2, iSMART and tcrdist(3) (12–15). Along with clusTCR, these are naive in the sense that they do not require epitope-specific or any other additional information about the sequences. We illustrate that clusTCR offers a 50 × performance increase (at 1 million sequences) compared to GLIPH2, which was found to be the second fastest clustering algorithm (fig. 3, table S1). In this study, it was illustrated that no single algorithm outperformed all others in terms of clustering quality. Therefore, we propose clusTCR as the most favourable option for clustering medium (> 200,000 sequences) to large (> 1,000,000 sequences) sized RepSeq data sets. Additionally, clusTCR provides the unique functionality of batch clustering, which allows processing of extremely large data sets, even if they do not fit into RAM. Finally, because clusTCR implements multiprocessing, it is perfectly suitable for high-performance computing infrastructures.

Along with clustering functionality, clusTCR provides tools for downstream analysis of the clustering results. These include calculation of cluster features such as cluster entropy, physicochemical properties and generation probability. Cluster features are particularly useful for downstream machine learning applications for TCR repertoire data. To this end, clusTCR may serve as an efficient tool for generating lower dimensional representations of highly complex immunosequencing data, while retaining the bulk of information contained in the original data set. clusTCR provides a classification model for evaluating the quality of the obtained clustering results. This model may serve as an *t* additional tool for lowering the dimensionality of the input data set. We found that *P_gen_* was the most important feature towards predicting clustering quality. Indeed, it has been illustrated previously that *P_gen_* is an important determinant for TCRs to recognize the same epitope (16). Although our model already achieves a solid performance (average AUC = 0.84 ± 0.07), we expect it to perform better when fitted with more data. Given the growing interest in TCR analysis, more ground truth data (i.e. epitope-labeled) is likely to be generated in the future.

## Conclusion

TCR clustering approaches are limited in their ability to scale up to the often extremely large immunosequencing data sets generated today (4, 17). clusTCR is a novel sequence-based CDR3 clustering approach, specifically developed to account for this limitation. We have illustrated that our approach is comparable to the current state-of-the-art in terms of clustering accuracy. At the same time, clusTCR provides a drastic speed improvement of > 50× over existing alternatives and allows clustering of large data sets that do not fit into RAM. To this end, this novel approach may unlock simultaneous processing of large RepSeq data sets, providing an effective tool for searching enriched groups of CDR3 sequences within a set of TCR repertoires.

## Bibliography

1. Mark M Davis, John D Altman, and Evan W Newell. Interrogating the repertoire: broadening the scope of peptide–mhc multimer analysis. Nature Reviews Immunology, 11(8):551–558, 2011.

2. Pieter Meysman, Nicolas De Neuter, Sofie Gielis, Danh Bui Thi, Benson Ogunjimi, and Kris Laukens. On the viability of unsupervised t-cell receptor sequence clustering for epitope preference. Bioinformatics, 35(9):1461–1468, 2019.

3. Mikhail Shugay, Dmitriy V Bagaev, Ivan V Zvyagin, Renske M Vroomans, Jeremy Chase Crawford, Garry Dolton, Ekaterina A Komech, Anastasiya L Sycheva, Anna E Koneva, Evgeniy S Egorov, et al. Vdjdb: a curated database of t-cell receptor sequences with known antigen specificity. Nucleic acids research, 46(D1):D419–D427, 2018.

4. Ryan O Emerson, William S DeWitt, Marissa Vignali, Jenna Gravley, Joyce K Hu, Edward J Osborne, Cindy Desmarais, Mark Klinger, Christopher S Carlson, John A Hansen, et al. Immunosequencing identifies signatures of cytomegalovirus exposure history and hla-mediated effects on the t cell repertoire. Nature genetics, 49(5):659–665, 2017.

5. Jeff Johnson, Matthijs Douze, and Hervé Jégou. Billion-scale similarity search with gpus. IEEE Transactions on Big Data, 2019.

6. Stijn Marinus Van Dongen. Graph clustering by flow simulation. PhD thesis, 2000.

7. Thomas D Schneider, Gary D Stormo, Larry Gold, and Andrzej Ehrenfeucht. Information content of binding sites on nucleotide sequences. Journal of molecular biology, 188(3): 415–431, 1986.

8. Zachary Sethna, Yuval Elhanati, Curtis G Callan Jr, Aleksandra M Walczak, and Thierry Mora. Olga: fast computation of generation probabilities of b-and t-cell receptor amino acid sequences and motifs. Bioinformatics, 35(17):2974–2981, 2019.

9. Sven Hellberg, Michael Sjoestroem, Bert Skagerberg, and Svante Wold. Peptide quantitative structure-activity relationships, a multivariate approach. Journal of medicinal chemistry, 30(7):1126–1135, 1987.

10. Asaf Madi, Asaf Poran, Eric Shifrut, Shlomit Reich-Zeliger, Erez Greenstein, Irena Zaretsky, Tomer Arnon, Francois Van Laethem, Alfred Singer, Jinghua Lu, et al. T cell receptor repertoires of mice and humans are clustered in similarity networks around conserved public cdr3 sequences. Elife, 6:e22057, 2017.

11. Anton J Enright, Stijn Van Dongen, and Christos A Ouzounis. An efficient algorithm for large-scale detection of protein families. Nucleic acids research, 30(7):1575–1584, 2002.

12. Hongyi Zhang, Longchao Liu, Jian Zhang, Jiahui Chen, Jianfeng Ye, Sachet Shukla, Jian Qiao, Xiaowei Zhan, Hao Chen, Catherine J Wu, et al. Investigation of antigen-specifict-cell receptor clusters in human cancers. Clinical Cancer Research, 26(6):1359–1371, 2020.

13. Huang Huang, Chunlin Wang, Florian Rubelt, Thomas J Scriba, and Mark M Davis. Analyzing the mycobacterium tuberculosis immune response by t-cell receptor clustering with gliph2 and genome-wide antigen screening. Nature Biotechnology, pages 1–9, 2020.

14. Koshlan Mayer-Blackwell, Stefan Schattgen, Liel Cohen-Lavi, Jeremy Chase Crawford, Aisha Souquette, Jessica A Gaevert, Tomer Hertz, Paul G Thomas, Philip Bradley, and Andrew Fiore-Gartland. Tcr meta-clonotypes for biomarker discovery with tcrdist3: quantification of public, hla-restricted tcr biomarkers of sars-cov-2 infection. bioRxiv, 2020.

15. Jacob Glanville, Huang Huang, Allison Nau, Olivia Hatton, Lisa E Wagar, Florian Rubelt, Xuhuai Ji, Arnold Han, Sheri M Krams, Christina Pettus, et al. Identifying specificity groups in the t cell receptor repertoire. Nature, 547(7661):94–98, 2017.

16. Mikhail V Pogorelyy, Anastasia A Minervina, Mikhail Shugay, Dmitriy M Chudakov, Yuri B Lebedev, Thierry Mora, and Aleksandra M Walczak. Detecting t cell receptors involved in immune responses from single repertoire snapshots. PLoS Biology, 17(6):e3000314, 2019.

17. Jennifer N Dines, Thomas J Manley, Emily Svejnoha, Heidi M Simmons, Ruth Taniguchi, Mark Klinger, Lance Baldo, and Harlan Robins. The immunerace study: A prospective multicohort study of immune response action to covid-19 events with the immunecode™ open access database. medRxiv, 2020.

