## Supplementary figures for "clusTCR: a Python interface for rapid clustering of large sets of CDR3 sequences"

Sebastiaan Valkiers, Max Van Houcke,
Kris Laukens & Pieter Meysman

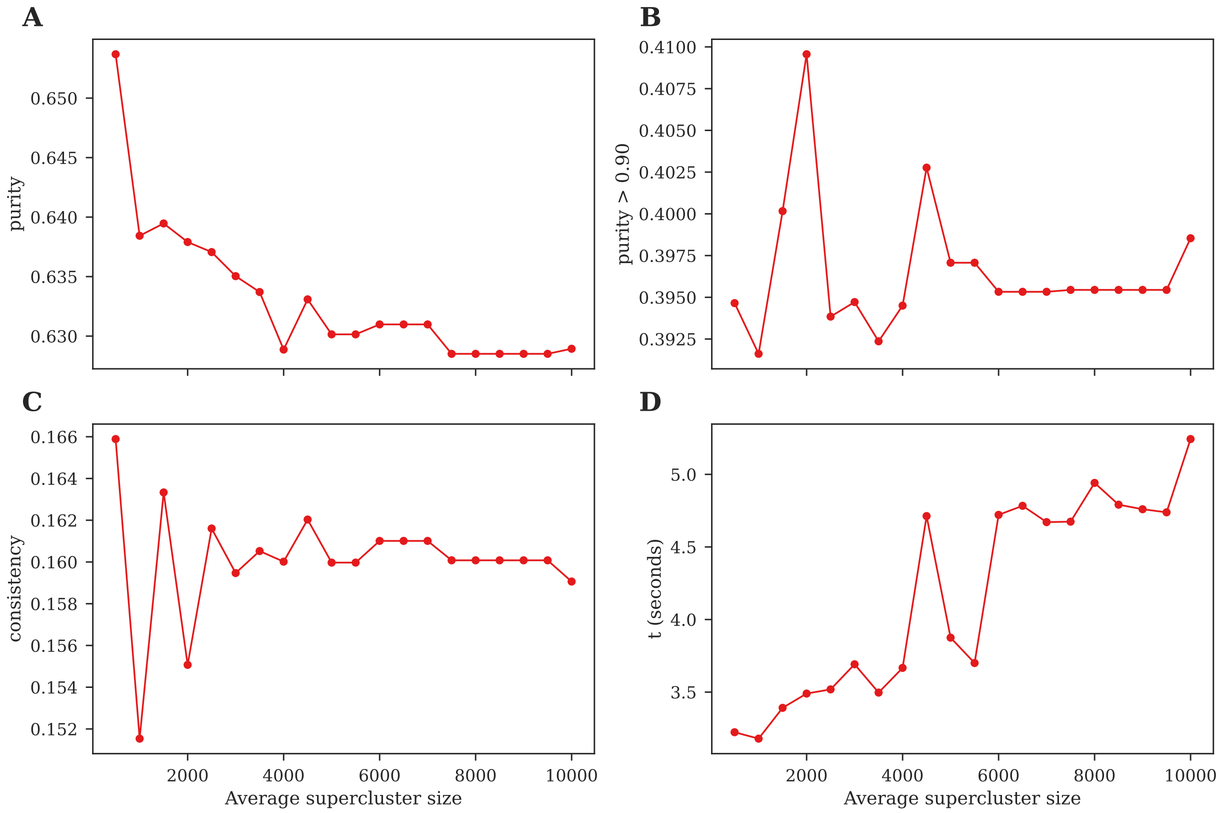


Figure S1: Influence of supercluster size on clustering performance on various clustering metrics, including **A.** purity. **B.** fraction of clusters with purity $> 90\%$, **C.** consistency, and **D.** the runtime of the algorithm.

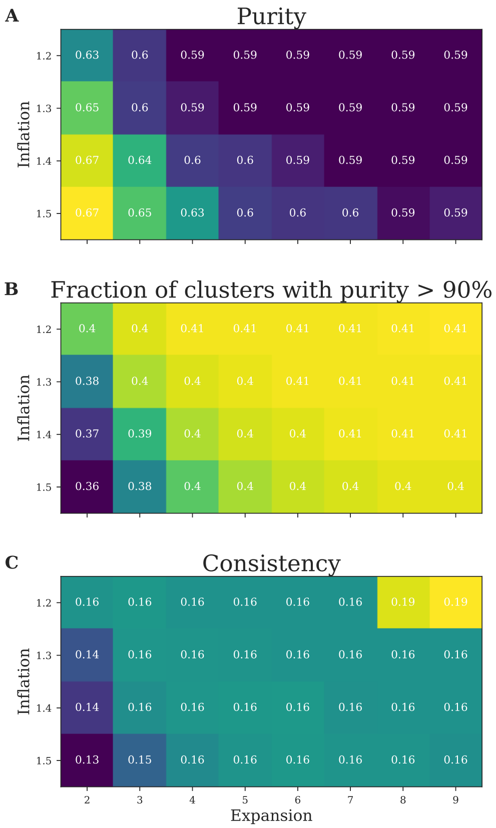


Figure S2: Evaluation of MCL hyperparameters. Different MCL hyperparameters were used and the clustering quality was evaluated using different evaluation metrics: **A.** purity, **B.** fraction of clusters with purity > 90% and **C.** consistency. A trade-off is observed between purity and consistency. *clusTCR* applies a default inflation of 1.2 and expansion of 2. This may not provide the optimal clustering results, but increasing these parameters results in decrease in speed of the algorithm.

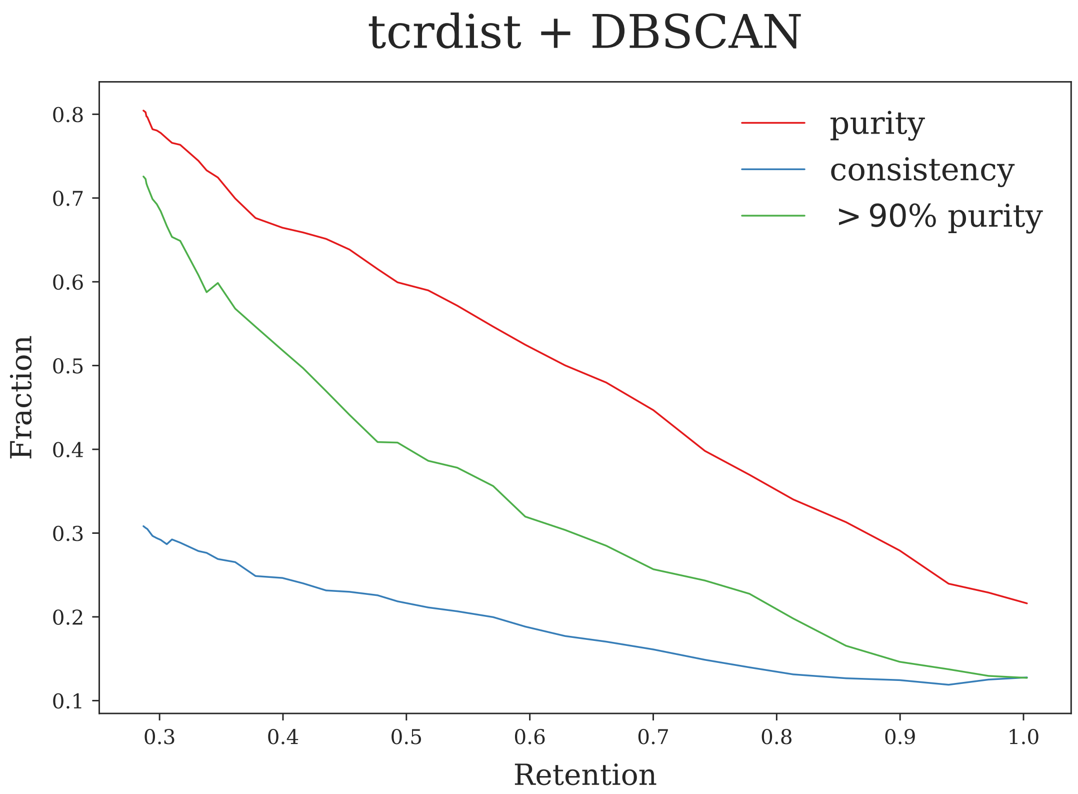


Figure S3: Performance of tcrdist + DBSCAN clustering. *tcrdist3* was used to calculate pairwise distances between CDR3 sequences. DBSCAN was used for clustering the sequences based on the pairwise distance matrix. By varying the DBSCAN similarity threshold (*eps* parameter), we evaluated the trade-off between retention, purity and consistency.

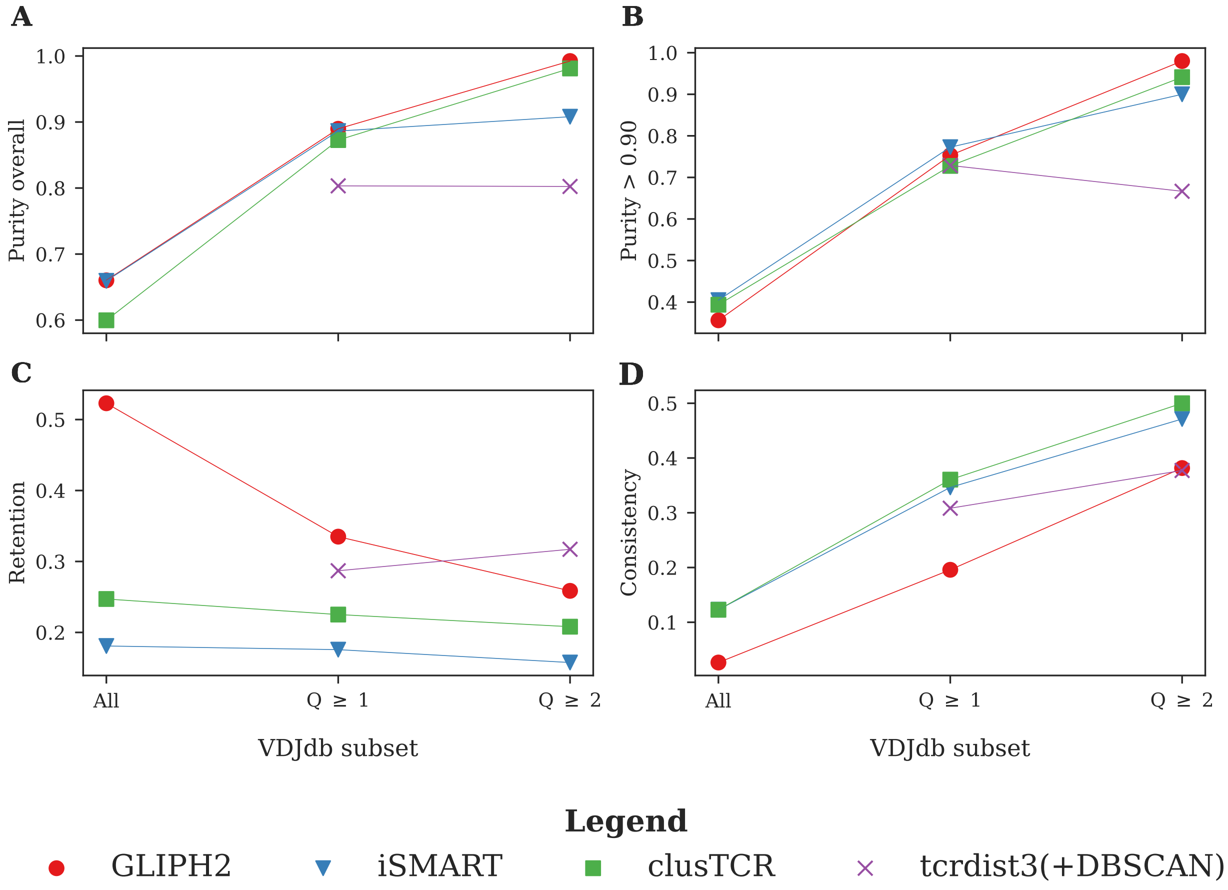


Figure S4: Clustering quality metrics of different TCR/CDR3 clustering approaches performed on different subsets of the VDJdb. **A.** Purity. **B.** Percentage of clusters with purity >90%. **C.** Retention. **D.** Consistency. Note that tcrdist3 was only used to calculate pairwise distances between the sequences, and DBSCAN was used to cluster them. Regular pairwise distance calculations were only possible for the two smaller subsets of VDJdb ($Q \geq 1$, $Q \geq 2$).

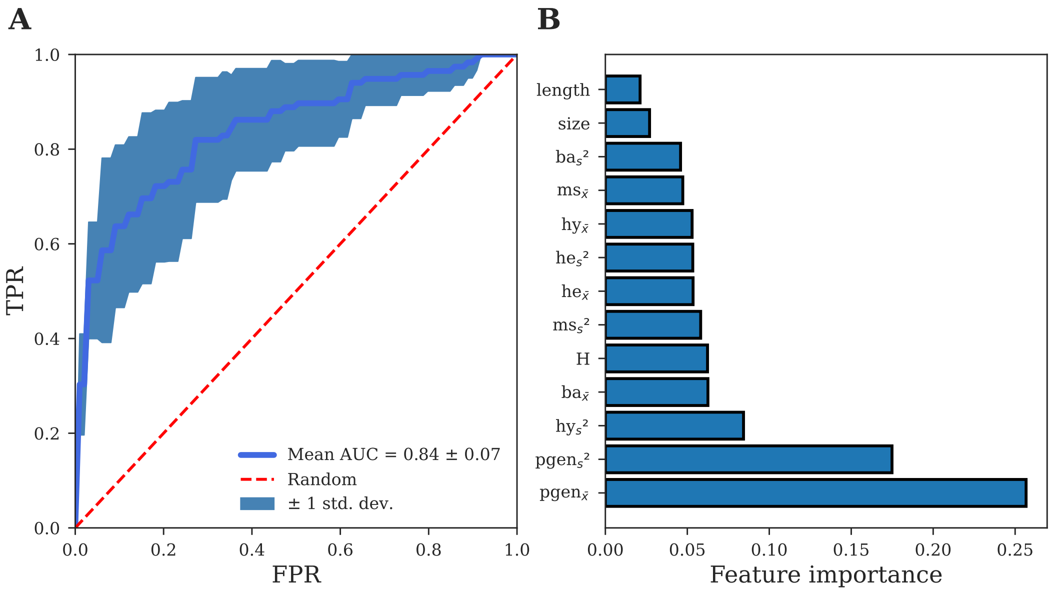


Figure S5: **A.** Cluster quality prediction model ROC curve. A classification model was built that identifies a good cluster from a bad cluster. Good clusters were defined as those clusters with purity $> 0.90$. The model was evaluated through 10-fold stratified cross-validation. The model had an average area under the curve (AUC) of 0.84, with a standard deviation of 0.07. This model is provided as the default model for cluster quality prediction in clusTCR. **B.** Importance of different cluster features in predicting cluster quality. Abbreviations: ba, basicity; he, helicity; hy, hydrophobicity; ms, mutation stability; pgen, generation probability.
