## Supplementary tables for "clusTCR: a Python interface for rapid clustering of large sets of CDR3 sequences"

Sebastiaan Valkiers, Max Van Houcke,
Kris Laukens & Pieter Meysman

Table S1: Runtime (in seconds) of different TCR/CDR3 clustering algorithms at different data set sizes (n = 3). * The runtime of tcrdist3 indicates pairwise distance calculation only, no clustering. Only one replicate was collected for 200K input sequences with tcrdist3.

| n_sequences ($\times 10^5$) | GLIPH2 | iSMART | clusTCR | tcrdist3* |
|---|---|---|---|---|
| 0.05 | $5.28 \pm 1.91$ | $2.14 \pm 0.18$ | $0.81 \pm 0.16$ | $19.84 \pm 0.73$ |
| 0.10 | $9.88 \pm 0.77$ | $5.68 \pm 0.4$ | $1.16 \pm 0.19$ | $73.11 \pm 1.91$ |
| 0.25 | $42.78 \pm 2.21$ | $24.79 \pm 1.06$ | $2.51 \pm 0.44$ | $439.91 \pm 17.58$ |
| 0.50 | $113.47 \pm 7.64$ | $92.7 \pm 3.74$ | $4.91 \pm 0.25$ | $1690.65 \pm 64.39$ |
| 1.00 | $307.86 \pm 19.44$ | $347.91 \pm 23.21$ | $13.32 \pm 1.17$ | $6450.21 \pm 64.19$ |
| 2.00 | $937.74 \pm 97.32$ | $1298.96 \pm 326.05$ | $32.96 \pm 7.28$ | 25342.9 |
| 5.00 | $4664.02 \pm 598.57$ | / | $117.99 \pm 27.74$ | / |
| 7.50 | $6833.25 \pm 310.21$ | / | $188.4 \pm 35.41$ | / |
| 10.00 | $9812.83 \pm 660.06$ | / | $239.53 \pm 30.31$ | / |
